# Self-compassion is linked to positive affect and buffers declines in self-esteem in low performers

**DOI:** 10.1101/2025.11.11.687618

**Authors:** Jan F Weis, Sören Krach, Frieder M Paulus, David S Stolz

## Abstract

Successes and failures shape affective experience and self-esteem. Self-compassion has been proposed as a protective factor, allowing individuals to acknowledge both strengths and shortcomings without excessive self-criticism. However, the mechanisms through which self-compassion influences changes in affect and self-esteem remain poorly understood. Here, we experimentally tested whether self-compassion modulates the links between performance feedback, affective experience, and self-esteem. Participants completed an effortful performance task and received trial-by-trial feedback while repeatedly rating their positive affect. Results show that self-compassion buffered against declines in self-esteem among poorly performing individuals, predicted higher overall positive affect throughout the task, and was associated with increased post-task self-esteem. Moreover, performance feedback predicted positive affect, which in turn predicted post-task self-esteem, although these pathways were not moderated by self-compassion. Together, these findings add to the growing evidence for how self-compassion impacts positive affect and self-esteem and may inform treatment strategies for clinical populations characterized by low self-esteem or heightened self-criticism.

## Introduction

Any action we take can fail or succeed. Our subsequent affect and resulting self-esteem depend on how we evaluate our successes and failures. While a particular response style towards such outcomes, self-compassion (Neff, 2003), has established implications for motivation, coping with stress, job satisfaction, or mental well-being (Breines & Chen, 2012; Galanakis et al., 2016; Kotera & Van Gordon, 2021; Neff, 2011; Neff et al., 2005), few laboratory experimental studies have tested how self-compassion modulates the impact of success and failure on self-esteem or positive affect (Waring & Kelly, 2019). Previous work has established that predictions of randomly occurring rewards, associated reward prediction errors, and even unrewarded effects of actions shape positive affect (Blain & Rutledge, 2020; Chew et al., 2021; Rutledge et al., 2014), but has neither assessed self-esteem or self-compassion. Further, although much of human behavior occurs independently of direct social comparisons, existing laboratory-based work on these processes has largely focused on socially comparative performance feedback (Czekalla et al., 2020, 2024; Müller-Pinzler et al., 2022, 2019; Schröder et al., 2025; Will et al., 2017; Williams & DeSteno, 2008; Zell & Strickhouser, 2020) and less on the impact of action outcomes (Stolz et al., 2020). The present study tries to provide these missing links by combining a novel, performance-based experimental task with computational modeling and thereby test if and how self-compassion shapes the affective and self-evaluative consequences of observing the outcomes of one’s own actions.

Self-esteem reflects the evaluation of oneself as a person, is often described as the feeling of being “good enough” (Orth & Robins, 2014), and is among the most frequently investigated topics in psychology (Muris & Otgaar, 2023). A large body of cross-sectional studies (Robins et al., 2002; Wagner et al., 2014) show that individuals with high (low) self-esteem hold generally positive (respectively negative) beliefs about themselves, including beliefs about abilities or skills, appearance, and social acceptance by others (Crocker & Park, 2004; Heatherton & Polivy, 1991; M. R. Leary & Baumeister, 2000; Tafarodi & Swann, 2001). High self-esteem individuals report higher self-efficacy, show more persistence in challenging tasks, and higher expectations regarding performance and success (Dutton & Brown, 1997; Müller-Pinzler et al., 2022; Zamfir & Dayan, 2022), even at objectively similar performance as individuals with lower self-esteem (Rouault et al., 2022). Importantly, links between self-esteem and beliefs about performance are likely bi-directional (Bentall et al., 2001; Seow et al., 2021): while high self-esteem is linked to success expectations, positive performance experiences also promote high self-esteem (Katyal et al., 2025; Müller-Pinzler et al., 2019; Neff et al., 2005; Rouault et al., 2019; Zamfir & Dayan, 2022).

A common and highly effective experimental method for influencing self-esteem is to present social evaluative feedback to participants (Eisenberger et al., 2011; Koban et al., 2017). For instance, inclusion or exclusion from group games (Zadro et al., 2004), or receiving surprisingly good (or bad) evaluations of one’s personality (Will et al., 2020, 2017) are significant drivers of self-esteem, in line with a role in tracking one’s social status (M. R. Leary & Baumeister, 2000). Here, self-esteem is especially responsive to the cumulative impact of recent prediction errors between the actual and predicted social feedback, rather than the overall valence of the feedback (Low et al., 2022; Will et al., 2020). Providing socially comparative performance feedback in cognitive tasks is another relevant influence, where feedback that is better than expected entails increases self-esteem or associated beliefs about own abilities, while feedback that is worse than expected has contrary effects (Müller-Pinzler et al., 2022, 2019; Rosi et al., 2019). For these reasons, it is thought that momentary self-esteem functions as an integrator of novel information potentially relevant to one’s current social status, which hinges on such factors as one’s social relationships and abilities (Shariff & Tracy, 2009).

Psychological theory holds that in close association with self-esteem, performance information also drives instantaneous affective experience, especially in response to one’s own actions (Tracy et al., 2009; Tracy & Robins, 2007). Accordingly, higher average levels of happiness and pride are positively linked to self-esteem (Stanculescu, 2012; Tracy & Robins, 2007), and individuals who experience low embarrassment or high pride during performance-based tasks update their ability-beliefs more positively in response to feedback (Müller-Pinzler et al., 2022). Similarly, receiving unsatisfactory feedback in response to one’s actions or inactions may elicit upward counterfactual thinking, a cognitive process about how alternative options could have produced better results (Gilovich & Medvec, 1995; Giorgetta et al., 2013). Aside from promoting learning and behavioral change, upward counterfactual thinking is also associated with the experience of regret, as supposedly reflected in the experience of lower positive affect (Gilovich & Medvec, 1995). These affective dynamics should thus arise in particular when available evidence favoring a certain course of action is ignored, and one’s behavior consequently entails suboptimal outcomes. Notably, the experience of regret has been associated with lower self-esteem in the long run (Righetti & Visserman, 2018). In summary, when trying to understand the impact of success and failure more fully, it is important to consider both instantaneous affective responses in ongoing tasks as well as changes in self-esteem.

As seen above, feedback on performance can impact individuals’ momentary affect and self-esteem. Notably, identical performance may lead to different affective experience and self-evaluation in different individuals. A personality trait that modulates these dynamics is self-compassion. Self-compassion describes an indulgent and kind approach to oneself in moments of self-inflicted suffering e.g. following a personal misstep (Neff, 2011). Being self-compassionate is thought to be beneficial for mental health and is linked to enhanced life satisfaction, happiness, optimism, self-efficacy, and a diminished experience of negative emotions (Liao et al., 2021; Muris & Otgaar, 2023; Neff, 2011). Additionally, individuals with higher self-compassion exhibit reduced self-criticism, stress, rumination, depression, anxiety, overall psychopathology, and neuroticism (Amirazodi & Amirazodi, 2011; Blackie & Kocovski, 2019; Han & Kim, 2023; Holas et al., 2023; MacBeth & Gumley, 2012; Muris & Otgaar, 2023; Neff, 2023; Neff & Vonk, 2009). So far, only few laboratory-based experimental studies have examined the role of self-compassion in self-esteem and affective experience. Empirical findings have demonstrated that self-compassion modulates affective responses to social feedback (M. Leary et al., 2007; Zhang et al., 2024). Given these insights, it appears likely that self-compassion shapes momentary affective responses to success and failure and, more globally, the impact of task performance on changes in momentary self-esteem.

In the present study, we used computational modeling to examine how performance information shapes affective experiences and self-esteem in dependence of self-compassion. Participants experienced trial-wise successes and failures in a novel, performance-based experimental paradigm – the *Learning-of-Effort-Efficacy* (LOEE) task. They additionally provided repeated ratings of their affective experience, indicated their self-esteem both before and after the task, and reported on their self-compassion after task completion. We examined how trial-wise and aggregate performance information predict affective experience and self-esteem, respectively, in dependence on individual levels of self-compassion. More specifically, we assessed how self-compassion modulates the impact of computationally-derived reward expectations, reward prediction errors, and ignored evidence on affective experience, and similarly, the impact of average task performance and positive affect on changes of self-esteem. We hypothesized that performance in the LOEE-task would predict changes in positive affect and in self-esteem, with a protective role of self-compassion in the face of low performance feedback.

## Methods

### Participants

We collected 85 complete datasets in our study. Of these, 11 datasets were excluded due to problems with calibrating the hand-dynamometer (in the following: power grip, see below) (*n*=4), or because participants showed extremely low variance in pride/happiness ratings (*n*=5) or effort expenditure (*n*=1). Last, one dataset was excluded because the participant accidentally took part twice (here, the person’s first dataset was used). For our final analysis we thus used data of *n*=74 participants (female=81%, age: *mean*=22.7, *sd*=3.7). Participants were students recruited mostly from the University of Lübeck or University of Applied Sciences Lübeck and gave written informed consent before starting participation. The study protocol was approved by the local ethics committee of the University of Lübeck (AZ 18-014).

### Procedure

After providing written informed consent, participants were seated in front of a desktop computer and completed a set of questionnaires online (Leiner, 2024). Afterwards, the power grip was calibrated and participants received detailed instructions about the LOEE-task on a computer screen. They performed two practice trials to verify the correct understanding of the task. After completion of the practice trials, the main task was started. During questionnaires, the calibration, instructions, and the LOEE-task, participants were seated alone in the laboratory in front of a desktop computer. In between each step, the experimenter joined the participants to answer potential questions about the procedures and the task before it started. After finishing the LOEE-task, participants again completed a set of online questionnaires, were debriefed and received a monetary compensation or partial course credit for participation in the study.

### Calibration of hand-dynamometer

Prior to the main task, a mini-game was conducted to calibrate the power grip, i.e., to assess the participant’s maximum voluntary contraction (MVC), which was necessary for scaling the feedback in the LOEE task (see below). To measure MVC, participants took the power grip into their dominant hand and were asked to hold it with the same hand for the remainder of the experiment. The mini-game was designed to incentivize participants to press the power grip as strongly as possible. Here, participants earned points if they exerted enough effort to reach a certain threshold. Their effort was visualized as the image of a blue flash which continuously grew or shrunk in size in relation to the amount of effort invested by the participant. The threshold was visualized as the white silhouette of a flash, and was reached if the blue flash filled that silhouette completely. If the participant exceeded the threshold, a star appeared as a reward and the required effort for the threshold of the next trial (i.e., the size of the silhouette) was increased (or lowered) by 6.75% with a probability of .67 (.33, respectively). If the threshold was not reached, a star marked with a red cross was shown, signaling non-attainment and the required effort for the threshold of the next trial was lowered (increased) by 6.75% with a probability of .67 (.33, respectively). We are confident that this procedure effectively assessed MVC, since all participants failed to reach the required threshold multiple times during all 15 calibration trials (*min*=3, *mean*=10.30, *sd*=2.30).

### The Learning of Effort-Efficacy (LOEE) task

#### Trial structure

In the LOEE-task, participants could earn money based on their performance while learning feedback contingencies in an uncertain environment. Each trial consisted of a choice phase, an effort execution phase, and a feedback phase. At the beginning of each trial, participants had five seconds to choose between a blue and a yellow button by clicking the arrow keys on the keyboard, using their non-dominant hand. After selecting a button, the image of a white flash appeared between the buttons, which indicated to participants that they could now invest effort by pressing the power grip. If the participant exerted at least 12.5% MVC within 1.75 seconds, the lightning turned blue, and the effort phase ended after 1.75 seconds. If 12.5% MVC was not reached by that time, the effort phase lasted until the threshold was reached. Next, participants received feedback on their performance of the given trial. If the participant had initially chosen the correct button (see below), they gained points in proportion to the effort they invested. They lost the same number of points if they had chosen the incorrect button. Since we calibrated the maximum effort for each participant prior to the experiment, gained or lost points were limited to −100 and 100 (corresponding to a maximum of 100% MVC). The feedback screen, shown for 4 seconds, displayed the correct button in this trial encircled in green and the incorrect button encircled in red. The number of points gained (or lost) were displayed simultaneously in green (or red, respectively). The points gathered in the entire task were later converted into a monetary payout of up to 7 Euros.

#### Input sequence

A predefined input sequence determined which button was correct at each trial of the task (see Fig. 1B), and the input sequence was the same for all participants. Participants were informed that in some periods of the task, one button would be correct with a higher probability and that these probabilities could change unpredictably. Also, they were informed that periods of stability in the probability could sometimes be rather long, and sometimes rather short. The precise frequency of probability changes was unknown to the participants. Thus, participants were required to track the winning contingencies of each button based on the feedback they received. Participants were informed that, across all trials, both buttons had the same probability of being correct.

**Figure 1.**
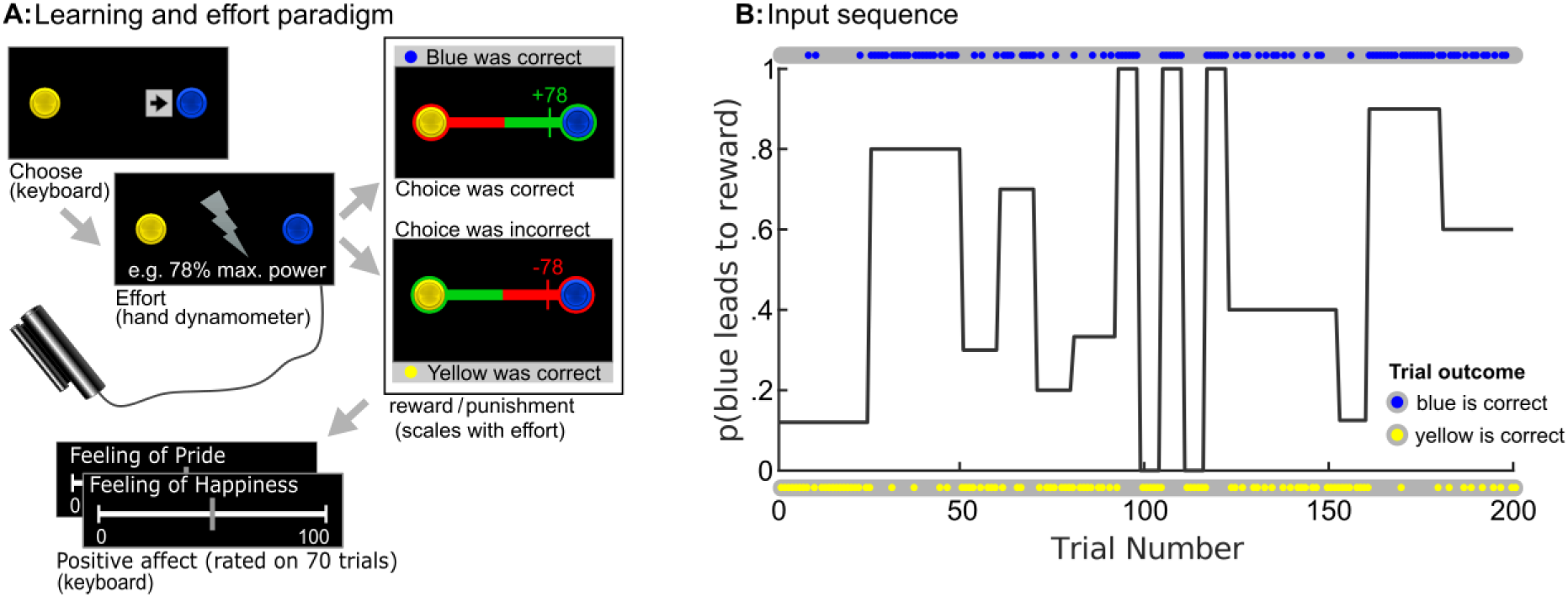
Experimental design **A)** On every trial, participants chose between two buttons. After their choice, they invested effort into their decision which determined how many points they won or lost given the eventual outcome. On every 2-3 trials, participants rated their positive affect. Participants were given no information about the winning probability of each button and had to track each button’s winning probability by updating their expectations based on previous observations. **B)** All participants received the same input sequence that determined which button was correct on each trial. The black line depicts the mean frequency with which “blue is correct” during predefined phases of the task. The outcome on every trial, i.e. which button was correct, is presented as blue or yellow dots.

#### Task related behavioral read-outs

Task related behavioral readouts covered affective experiences (happiness and pride) as well as effort and choice data. These were used to validate the experimental task behavior and test the hypotheses on determinants of self-esteem.

*<u>Choice data.</u>* Each trial participants had to select either the yellow or the blue button. If the participant’s selected button matched the outcome button, the choice was correct. The percentage of correct choices for each participant was calculated as the sum of correct choices divided by the number of trials. Similarly, if the participant chose the correct button in the previous trial and decided to choose the same button again, the trial was coded as a “stay trial”. The stay probability was calculated as the conditional probability of staying with their previous choice given the previous outcome.

*<u>Effort data.</u>* Participants were required to exert effort on every trial to progress in the experiment. The amount of exerted effort dependent on their prior knowledge about button contingencies acquired from previous trials. To test if participants actually used their acquired knowledge, we calculated a hypothetical feedback that they would have obtained if they had invested the same, average level of effort on all trials, i.e., irrespective of having chosen the correct or incorrect button. Put differently, this hypothetical feedback would arise if subjects did not adjust their effort investment on each trial in dependence of their current knowledge of the task.

*<u>Affect ratings.</u>* On every 2-3 trials after the feedback was displayed, participants were requested to sequentially rate their feeling of happiness and pride on two separate scales using a slider. When rating their happiness (or pride), participants were asked “How happy (proud) are you about the feedback of the last trial?” and should rate their feeling on a scale from “not happy (proud) at all” over “neutral” to “very happy (proud)”. Happiness and pride ratings were then read out as a number between 0 and 100. In total, 200 trials were performed with 70 trials in which pride and happiness were provided by each participant. 38 out of all 74 participants included in the final analysis always rated pride before happiness, and the rest the other way around. Since ratings of pride and happiness were highly correlated (within-subject Pearson r, *median*=.90, *IQR*=.17), we took their trial-wise average to compute momentary positive affect for each trial.

### Questionnaires

As a measure of self-esteem, participants completed the revised German version of the State Self-Esteem Scale (SES) (Rudolph et al., 2020). This scale measures momentary self-esteem, divided into three domains: social, performance and appearance self-esteem (Heatherton & Polivy, 1991). Participants completed the SES both before and after the task. Scores for each domain were calculated as means of all respective items. Questionnaire items exhibited high internal consistency with a Cronbach’s alpha of 0.85, similar to scores found in the literature (Heatherton & Polivy, 1991; Rudolph et al., 2020). In the remainder of this manuscript, unless specified otherwise, we use “self-esteem” to refer to the performance domain measured by the SES. Additionally, the Self-Compassion Scale (Neff, 2003) was completed by each participant after the task to evaluate their level of self-compassion. Again, self-compassion scores were presented as the mean of all items, exhibiting high internal consistency (Cronbach’s alpha of 0.9), which is consistent with previous research (Holas et al., 2023) (see Fig. S3 for descriptive statistics). A summary of the internal consistency estimates and correlations among the questionnaire measures is provided in the supplements (see Table S1).

### Statistical analysis

All analyses were performed using the R Statistical Software (R Core Team, 2024) in RStudio (RStudio: Integrated Development for R. RStudio, 2023) and MATLAB (*The MathWorks Inc*, 2022).

*Computational modeling of choice data*.

Previous work showed that momentary positive affect is driven by reward expectations and reward prediction errors (Blain & Rutledge, 2020). To derive these quantities and test their impact on positive affect in our data, we modeled trial-by-trial choice data using a Rescorla-Wagner (RW) model (Sutton & Barto, 2018) in the MATLAB-based VBA Toolbox (Daunizeau, 2025; Daunizeau et al., 2014). The model was specified by: Expectation_t_ = Expectation_t-1_ + α * (Outcome_t-1_ – Expectation_t-1_) where α represents the learning rate with which new information is integrated into prior beliefs and t is the trial number. Model inversion was run using the mixed-effects empirical Bayes approach implemented in the VBA_MFX function of the VBA toolbox. Fitting the RW-model to each participant’s choice data provided a good fit to the overall learning process (balanced classification accuracy: *mean*=0.71, *sd*=0.09; Fig. 3a)

Based on the individual RW-model fits, we then computed latent variables reflecting trial-wise performance information, which included reward expectancy (rewEXP), reward prediction error (rewPE)(Fig. 3b) and ignored evidence. Reward expectancy was specified as the difference of effort-dependent expected gain and expected loss on a given trial t, i.e. by rewEXP_t_ = p(correct|choice)_t_ * %MVC_t_ – p(incorrect|choice)_t_ * %MVC_t_. Reward prediction error was specified by rewPE_t_ = feedback_t_ – rewEXP_t_. Last, ignored evidence was calculated for trials on which a participant did not chose the more likely option and then lost (and set to 0 on all other trials). On these trials, ignored evidence was computed as p(correct)_t_ for the more likely but unchosen option, minus .5 (see Fig. 3c). Accordingly, we expected that participants who chose the less likely option contrary to the available evidence and subsequently lost would engage in upward counterfactual thinking and experience regret, reflected in reduced positive affect.

### Validation of task behavior

To assess the validity of our experimental design, we performed statistical tests using base R (R Core Team, 2024). Variables were tested for normality using the Shapiro-Wilk test. For variables where this test indicated a violation of normality assumptions (i.e. percentage correct choices, stay probability after correct), we used the Wilcoxon signed-rank test. Otherwise, parametric tests were performed using a simple one-sided t-test (i.e. average feedback, stay probability after incorrect) or a paired two-sided t-test (i.e. positive affect after outcome). Within-subject correlation values of positive affect and average feedback were normalized using the atanh()-function and subsequently tested against 0 using a one-sided t-test.

### Analyses of determinants of self-esteem and positive affect

To model trial-by-trial dynamics in positive affect, we ran a linear mixed-effects regression model using latent variables described above. Specifically, as first-level predictors, we used our latent performance-information variables (now referred to as performance information), including reward expectancy, reward prediction error and ignored evidence. Additionally, we included trial progression as a nuisance variable. As second-level predictors, we included pre-task self-esteem and self-compassion. We included a fixed intercept and fixed slopes for all predictors, as well as subject-wise random intercepts and slopes for trial progression and performance information to account for individual differences in feedback processing. Since self-compassion could moderate positive affective responses to feedback, we included cross-level interaction terms between self-compassion and performance information in our model (see Table S2 for more information regarding the development of the model). All first-level predictors were z-scaled within participants, and second-level predictors were centered across participants.

Mixed-effects regression models were run using the lmer()-function of the lme4-package (Bates et al., 2015), using the bobyqa-optimizer of the optimx-package (Nash, 2014; Nash & Varadhan, 2011). A null model was defined including only a fixed intercept and random intercepts for each subject. After sequentially adding fixed effects for our first-level predictors, we then incrementally added random effects for each first-level predictor. Last, second-level variables and their cross-level interactions with first-level predictors were added. Statistical tests involving marginal effects of mixed-effects models were performed using the emmeans- and marginaleffects packages (Arel-Bundock et al., 2024).

To examine whether the LOEE-task induced changes in self-esteem, we ran a mixed model with self-esteem as the dependent variable, time (Pre-Post), domain (performance vs. combined average of other domains) and their interaction as independent variables, using the lmer()-function. For the domain variable, the self-esteem ratings of the performance domain were contrasted with the combined and averaged ratings of the social and appearance domain.

Between-subject variability in post-task self-esteem was investigated by running linear regression models using the lm()-function included in base R. We first introduced the predictors (i.e. positive affect) into the model in a stepwise manner, and later introduced interaction terms with self-compassion. All predictors were centered between participants.

## Results

### Subjects use acquired knowledge to adjust their effort

Basic task behavior indicated that participants learned about the winning contingencies of the buttons and changed their behavior in response to correct or incorrect outcomes. Precisely, participants performed significantly better than chance (i.e., 50%) with 58.5% correct choices on average (Fig. 2A) (*median*=0.58, *IQR*=0.06, *W*=2576.5, *one-sided p*<0.001). Relatedly, average feedback across all trials was significantly higher than 0 (*mean*=8.20, *sd*=4.83, *t*(73)=14.6, *one-sided p*<0.001) (Fig 2B). When prompted to select a button, participants decided to repeat their previous choice significantly more often than chance (i.e., 50%) after a preceding success (*median*=0.84, *IQR* 0.19, *W*=2767, *p*<0.001) but significantly less often than chance after a preceding failure (*median*=0.42, *IQR* 0.21, *W*=610.5, *p*<0.001) (Fig. S1A).

**Figure 2.**
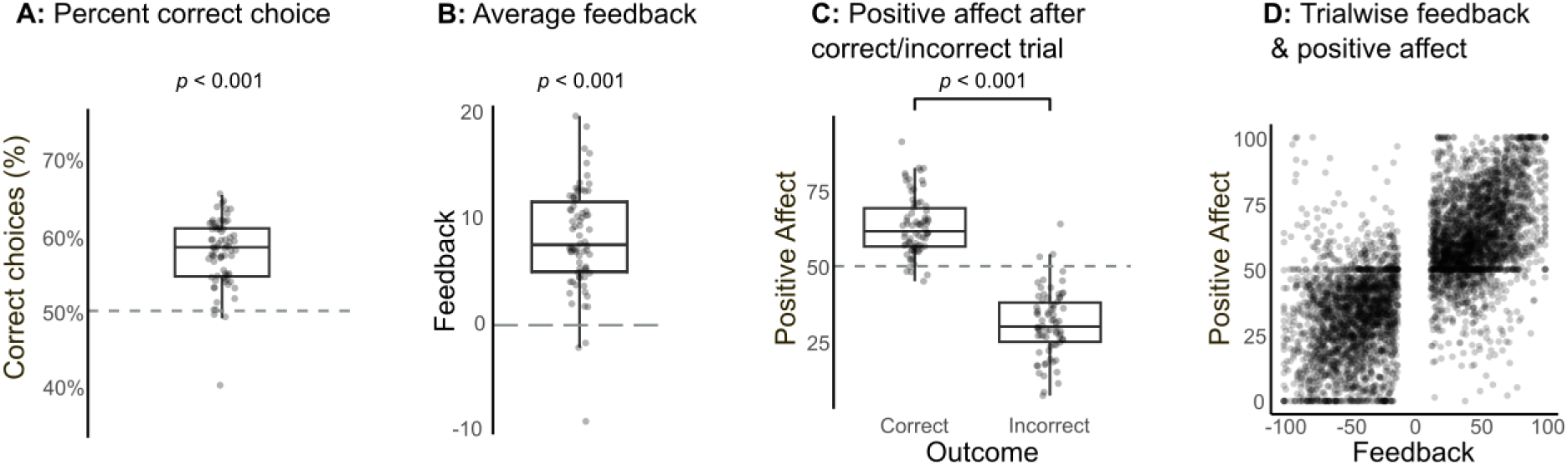
A) Participants performed significantly better than chance (50%). **B)** Participants gained significantly higher feedback than chance (0). **C-D)** Positive affect was higher after correct than incorrect trials and was positively correlated with the size of the previous feedback. Note that participants always had to show at least 12.5% MVC before the trial continued, leading to (absolute) feedback not being lower than 13 points.

**Figure 3.**
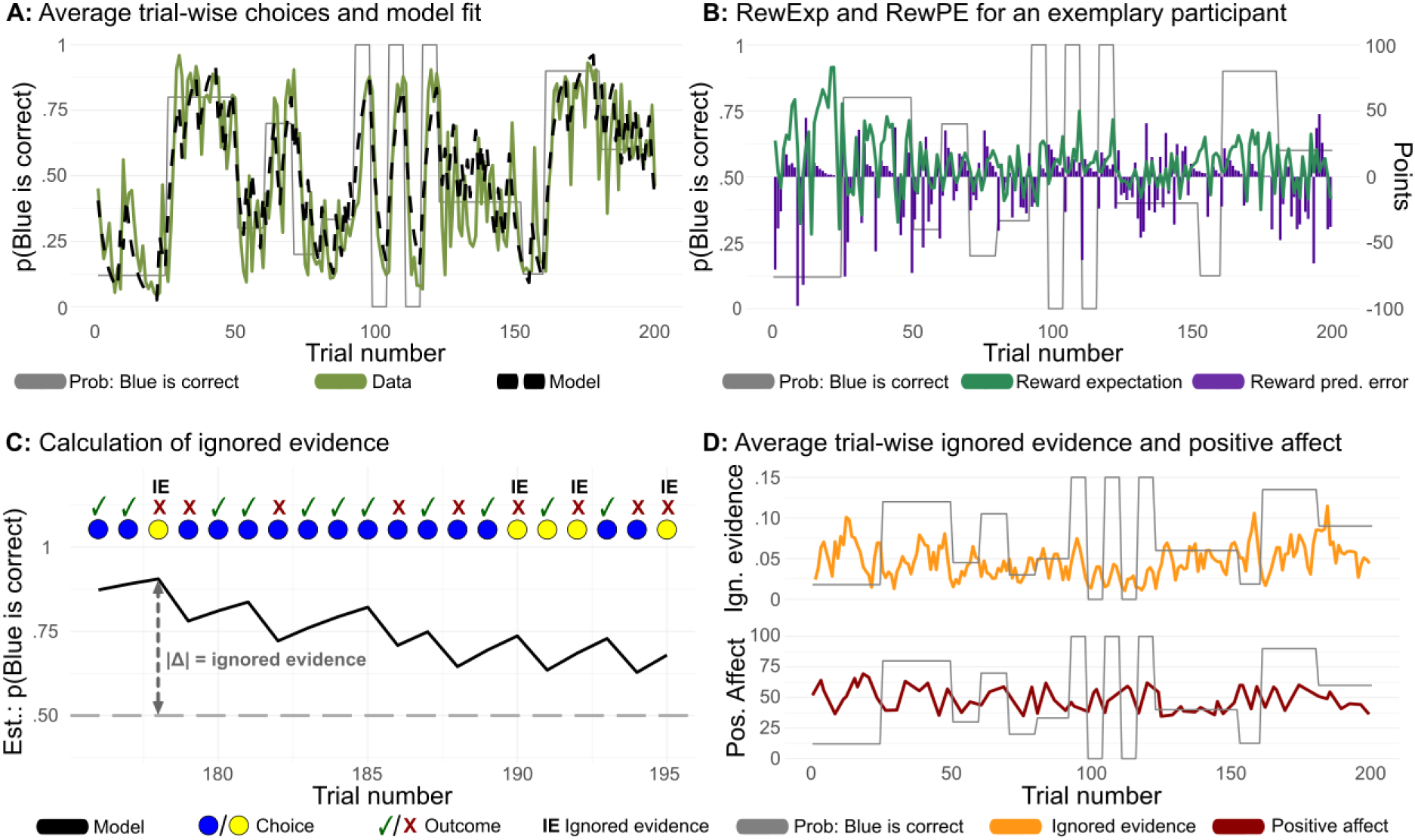
Rescorla-Wagner model accurately tracks participants’ choice. **A)** Plotted trajectories of the average choices, trial-by-trial, of all participants (light green) and the trial-by-trial estimation of the model across all participants (dashed black line), which approximately follow the actual winning probabilities for blue. Model’s choice closely resembles choice behavior of participants. **B)** Trajectories of the latent variables reward expectation (turquoise) and reward prediction error (violet) derived from the RW-model. **C)** Calculation of ignored evidence in a trial by taking the absolute difference between the model’s estimation for p(correct|choice) and .5. In a trial, ignored evidence was calculated when participants choice was incorrect and had a lower probability of winning given the model’s estimation (p(correct|choice)<.5). Higher values of ignored evidence serve as a proxy for greater levels of experienced regret. **D)** Plotted trajectories of ignored evidence (orange) and positive affect (dark red), trial-by-trial, across all participants. Trajectory of ignored evidence was smoothed with a moving average kernel size of 3.

Further, participants adjusted their effort depending on their expectation to have chosen the correct button. This was tested by comparing the participants’ actual average feedback against a hypothetical feedback that they would have obtained if they had invested the same, average level of effort on all trials (see methods). Here, we found that actual feedback significantly exceeded this hypothetical feedback (Fig. S1B) (*t*(73)=6.03, *p*<0.001; hypothetical feedback *mean*=7.32, *sd*=4.60). Together, this demonstrates that participants used their knowledge about the winning contingencies to guide effort investments, which allowed them to increase the payout in the task.

Regarding participants’ affective responses to trial outcomes, we found that participants reported significantly higher levels of positive affect in response to a correct trial (*median*=61.5, *IQR* 12.7) than an incorrect trial (*median*=30.4, *IQR* 13.0, *W*=5408, *p*<0.001) (Fig. 2C). In addition, we found a significant positive correlation between feedback size and subsequent positive affect (Fig. 2D) (Spearman correlation: *mean r*=0.73, *mean Fisher-Z-transformed r*=1.11, *t*(73)=24.49, *p*<0.001).

Collectively, these behavioral results suggest that participants understood the task, learned its structure, and guided their effort based on knowledge about the buttons, while also exhibiting expected changes in affective experience according to their performance.

### Self-compassion and performance information drive positive affect

The primary drivers of positive affect during the task were performance information (e.g., reward expectation) and self-compassion. We ran a linear mixed-effects model that explained 71% of the variance (*conditional R^2^*=0.71; *marginal R²*=0.52) with all first-level predictors reaching significance (see Fig. 4). Specifically, while trial progression (*β*=-4.76, *s.e.*=0.95, *p*<0.001) and ignored evidence (*β*=-1.51, *s.e.*=0.33, *p*<0.001) had negative effects, reward expectation (*β*=7.78, *s.e.*=0.50, *p*<0.001) and reward prediction errors (*β*=11.09, *s.e.*=0.72, *p*<0.001) had positive effects. These results demonstrate that positive affect was influenced by the anticipation of receiving a reward and when rewards exceeded these expectations, in line with previous findings (Rutledge et al., 2014). On the other hand, ignoring available evidence by choosing the less likely option and losing led to lower levels of positive affect, reflective of momentary regret.

**Figure 4.**
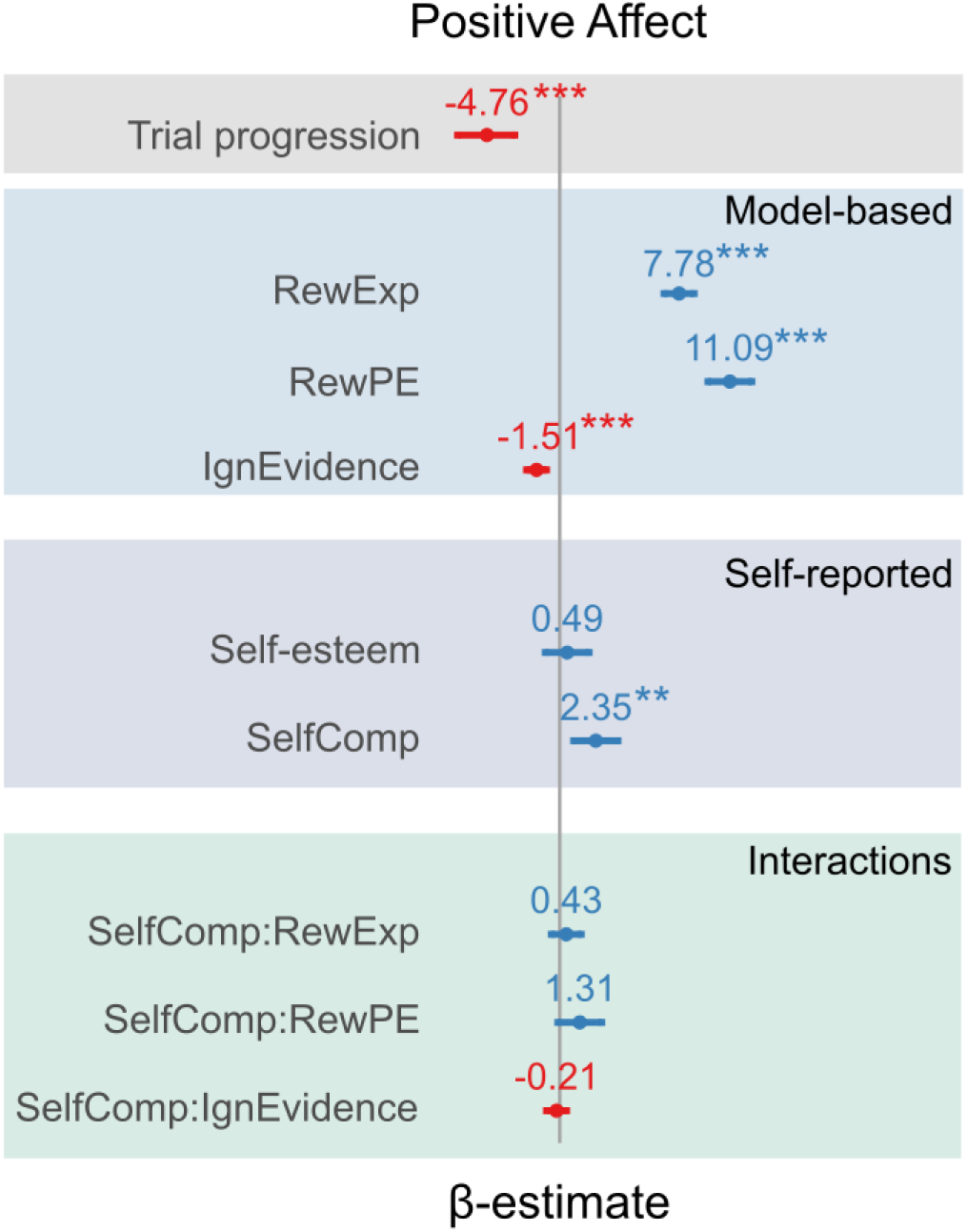
Performance information are the main driver of positive affect. Running a linear mixed-model using latent variables derived from the RW-model confirmed that performance information (reward expectation, reward prediction error and ignored evidence) along with self-compassion and trial progression best predict positive affect. ** *p* <.01, *** *p* <.001 RewExp = reward expectation; RewPE = reward prediction error; IgnEvidence = Ignored evidence; SelfComp = self-compassion;

For between subject variables, we found a significant effect for self-compassion (*β*=2.35, *s.e.*=0.73, *p*=0.002) but no effect for pre-task self-esteem (*β*=0.49, *s.e.*=0.73, *p*=0.51), similar to our behavioral model of trial-wise positive affect (see Fig. S2). Moreover, interactions between the model-based predictors and self-compassion remained non-significant (all *p*s>0.05).

### Self-esteem diminishes during task performance

As expected, due to the experimental context, participants’ performance-related self-esteem changed from before to after the task, while other domains of self-esteem remained constant (Fig 5). Using a mixed-model that explained 48% of the variance (*conditional R^2^*=0.48, *marginal R^2^*=0.09), we tested if self-esteem was predicted by time (Pre-Post), domain (performance vs. other domains) and their interaction. Across both timepoints, ratings of performance-related self-esteem were significantly higher compared to the other domains (*β*=0.15, *s.e.*=0.02, *p*<0.001). Moreover, we found a negative effect of time on self-esteem ratings (*β*=-0.07, *s.e.=*0.03, *p*=0.012), which, importantly, differed between domains. Specifically, self-esteem in the performance domain decreased significantly from before to after the task (*W*=1418.5, *paired p*<0.001), and this change was stronger than in the other self-esteem domains (planned contrast of the interaction of domain and time: *β*=-0.05, *s.e.*=0.02, *p*=0.012). As detailed next, we then aimed to map these changes in performance self-esteem onto overall task performance, affective experience, individual differences in self-compassion, and the interplay of these variables.

**Figure 5.**
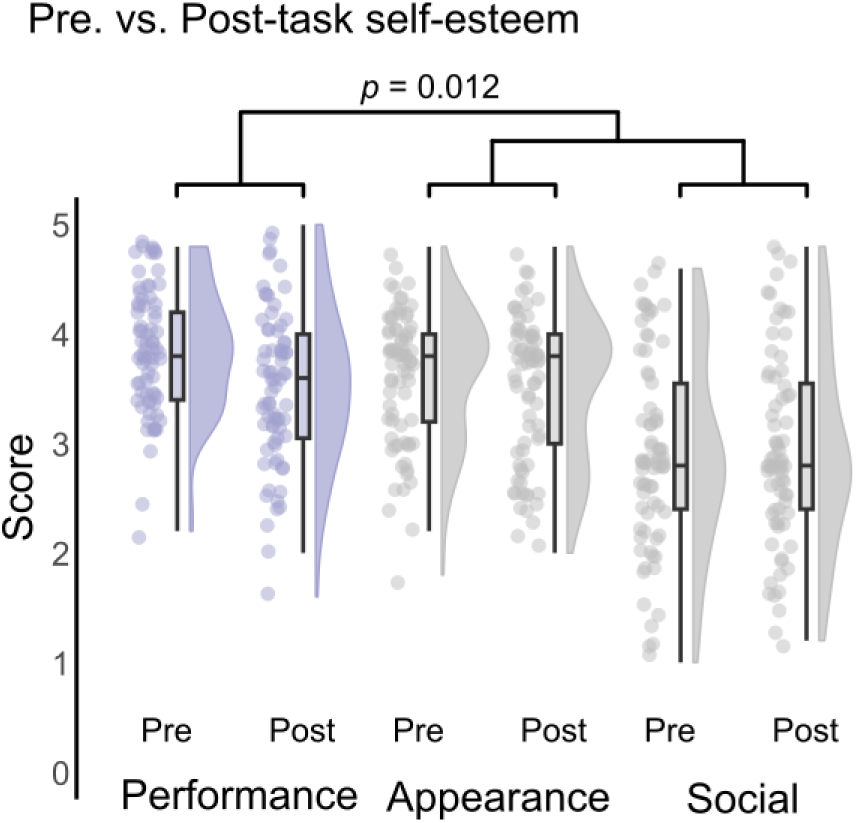
Reduction in performance self-esteem post task is significantly greater than self-esteem changes across social and appearance-related self-esteem domains.

### Self-compassion and positive affect explain post-task self-esteem

We found that positive affect and self-compassion predicted post-task self-esteem, and that self-compassion moderated the impact of performance feedback on post-task self-esteem. The model explained a significant amount of variance in post-task self-esteem with *R^2^*=0.59 (*F*(6,67)=18.8, *p*<0.001). As expected, pre-task self-esteem (*β*=0.36, *s.e.*=0.06, *p*<0.001) was a significant predictor of post-task self-esteem. Further, we found that average positive affect (*β*=0.14, *s.e.*=0.06, *p*=0.01) and self-compassion (*β*=0.14, *s.e.*=0.06, *p*=0.03) but not average feedback (*β*=0.05, s.e.=0.05, p=0.376) were predictive of higher levels of self-esteem after the task (Fig. 6a).

**Figure 6.**
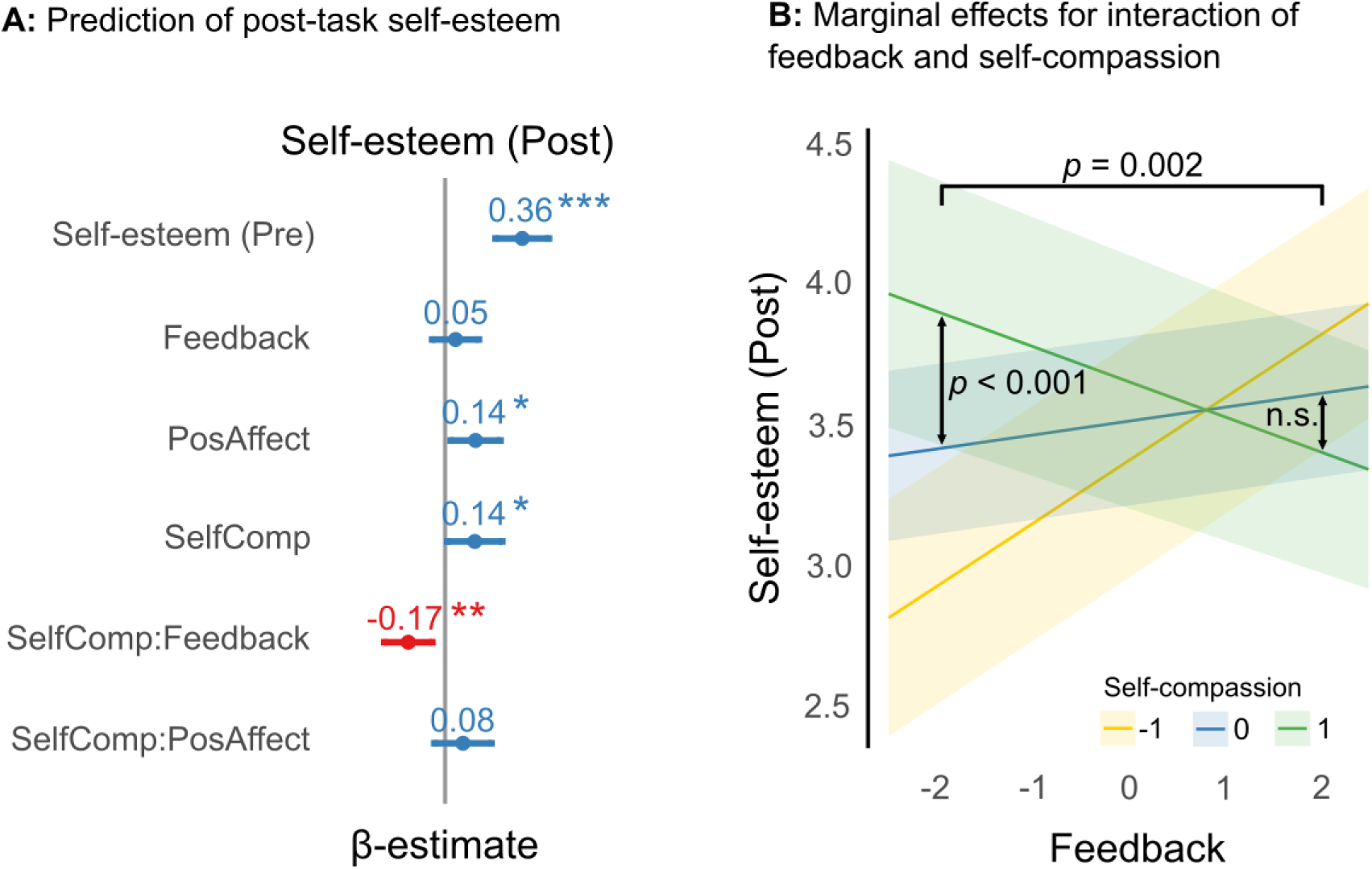
Self-compassion modulates the effect of overall performance feedback on self-esteem. **A)** A linear regression revealed self-esteem (Pre), positive affect, and self-compassion as significant predictors of self-esteem (Post). Further, self-compassion modulates the effect of feedback on self-esteem. **B)** Marginal effects of the interaction between self-compassion and feedback indicate that self-compassion is particularly effective in mitigating the effects of low feedback on self-esteem (see text for details). Feedback and self-compassion scores were z-standardized; a value of −1 indicates one standard deviation below the sample mean. * *p* < .05, ** *p* < .01, *** *p* < .001 PosAffect = positive affect; SelfComp = self-compassion; Pre = pre-task; Post = post-task

In line with our hypothesis, we found a significant interaction between self-compassion and average feedback (*β*=-0.17, *s.e.*=0.06, *p*=0.003), suggesting that higher levels of self-compassion impact how feedback affects post-task self-esteem. Exploring the marginal effects of this interaction (see Fig. 6b) revealed that for participants who received comparably less rewarding feedback, those with high self-compassion had higher post-task self-esteem than those with average or low self-compassion (*β*=0.49, *s.e.*=0.14, *p*<0.001). In contrast, for participants who received, on average, more rewarding feedback, no such effects were found (*β*=-0.21, *s.e.*=0.12, *p*=0.09). Moreover, the absolute difference in estimated post-task self-esteem between high and average self-compassion participants was larger for participants who received, on average, rather low feedback compared to those receiving rather high feedback (*β*=-0.35, *s.e.*=0.11, *p*=0.002). No significant interaction between self-compassion and average positive affect was found (*p*=0.21).

These results underline that the affective experience during an effortful task shapes post-task self-esteem in the performance domain above and beyond pre-existing differences in self-esteem. Similarly, self-compassion contributes to post-task self-esteem and, in addition, appears to buffer the effects of less rewarding feedback on self-esteem. However, no evidence was found to suggest that the affective experience during the task predicts self-esteem in dependence of individual differences in self-compassion.

## Discussion

Self-esteem and self-compassion are integral parts of an individual’s well-being, capturing both the evaluation of one’s worth as a person and the capacity to treat oneself with kindness. Despite its conceptual relevance, the role of self-compassion in shaping changes in self-esteem has not yet been explored in depth. In this study, we used computational modeling to show how self-compassion influences changes in self-esteem via affective experience and performance feedback. Using a task linking learning and effort execution, we find that self-compassion is particularly effective in reducing the adverse effects of low performance during task execution on self-esteem. Further, trial-wise performance information was associated with positive affect experienced during the task, which in turn predicted changes in self-esteem. Importantly, changes in self-esteem were predicted by subjective, rather than objective, performance evaluations, implying that self-compassion may promote outcome reappraisal and affective regulation.

Engaging in compassionate self-treatment has been proposed to foster adaptive self-evaluations (M. Leary et al., 2007; Neff, 2011). Allowing individuals to accept both strengths and weaknesses (or positive and negative feedback) as legitimate parts of the self (Crocker et al., 2006), could reduce the impact of external evaluations on self-esteem, and consequently reduce its fluctuations (Neff & Vonk, 2009). Consistent with this perspective, our study demonstrates that self-compassion modulated the impact of overall task feedback on self-esteem. As predicted, this effect was primarily driven by participants achieving relatively low levels of performance. Notably, individual differences in overall performance alone did not predict changes in self-esteem, indicating that participants’ self-esteem updates were shaped less by objective performance but rather by their subjective evaluation thereof. That is, when participants are asked to actively reflect on their abilities (e.g., during completion of the post-task self-esteem scale), self-compassion could help to downregulate the impact of recent personal failures, thereby reducing the effect of low performance on self-esteem. Evidence for this assumption comes from studies that demonstrate a connection between self-compassion and the use of cognitive reappraisal strategies, in particular reinterpreting and positively reconstructing events (Allen & Leary, 2010; Doorley et al., 2022).

To build on these initial insights, future experimental work should aim to establish whether practicing self-compassion does indeed causally elicit deliberate self-reflection (e.g. acknowledging the imperfection of human kind and the self) as a means of processing feedback and therefore helps preserve self-esteem (Brotzeller et al., 2025; Nowak et al., 2023). A further possibility that could be amenable to experimental investigation is that self-compassion does not reduce the overall intensity of affective reactions to feedback (especially negative reactions), but rather shortens their duration and mitigates maladaptive cognitive processes such as rumination and self-criticism (Blackie & Kocovski, 2019). Last, the stabilizing effect of self-compassion on self-esteem could in principle also arise from a broad attenuation of both positive and negative affective responses to feedback. This, however, seems unlikely in view of our findings.

Interestingly, the self-esteem of our participants decreased during the experiment, even though objective performance showed that the vast majority successfully learned the task structure and used their knowledge of that structure to optimize their payout. Furthermore, our task did not provide an external reference against which participants could compare their performance (like performance feedback relative to other participants). Together, this stresses the potential importance of subjective performance evaluations. Previous research demonstrated that self-esteem is sensitive to recent expectation violations, rather than the overall valence of the social feedback (Low et al., 2022; Will et al., 2017). Consequently, even objectively positive feedback may lower self-esteem if it falls short of one’s expectations. This view implies that participants approached our task with some implicit expectation of their overall performance, which was then violated, e.g. by the rather frequent negative feedback (42% on average). If true, this violation of performance expectations on the task-level might have reduced performance-related self-beliefs (Czekalla et al., 2024; Müller-Pinzler et al., 2022, 2019; Schröder et al., 2025), resulting in lower self-esteem. This would suggest that predicting relevant events and tracking violations of these predictions is a key driver of self-esteem dynamics in both social (Low et al., 2022; Will et al., 2017) and performance domains (e.g. expectations about social interactions/achievements in competitions, respectively). To establish whether such a potential unifying mechanism of self-esteem regulation indeed exists, a first relevant step would be to formally measure predictions and prediction violations in various domains within the same participants. Then, one could assess how strongly their impact on changes in affect and self-esteem corresponds between these domains.

Previous research has shown that highly self-compassionate individuals perceive neutral social feedback as more positive and experience less negative affect when evaluated by others (M. Leary et al., 2007; Zhang et al., 2024). In contrast, our findings showed that performance information predicted positive affect during the task, which in turn predicted self-esteem at the end of the task – while self-compassion did not modulate either of these effects. This suggests that self-compassion does not moderate immediate responses in positive affect to feedback nor does it modify the influence of positive affective experiences on self-beliefs. Still, higher self-compassion was linked to both greater positive affect and higher self-esteem suggesting that self-compassion may shape global self-beliefs through multiple pathways: directly, by fostering higher self-esteem, and indirectly, by enhancing positive affect, which in turn supports self-esteem. While not evident in our findings, we highlight that moderating effects of self-compassion may emerge primarily when assessing negative affective experiences (e.g. embarrassment, guilt, or shame) rather than the positive ones measured here. This would also be in line with the notion that self-compassion regulates affective and cognitive responses to personal failure, but not success (Neff, 2011). Even though positive and negative affect are inversely correlated, their frequency and intensity may vary independently (Diener et al., 1985), and they may even occur simultaneously (i.e. bittersweet moment) (Hoemann et al., 2017). Thus, testing the effects of self-compassion on negative affect remains an important research question, especially because affective experiences (e.g. pride, embarrassment) have been linked to the formation of performance-related self-beliefs (Czekalla et al., 2024; Müller-Pinzler et al., 2022). For instance, embarrassment has been shown to increase after negative performance feedback (Müller-Pinzler et al., 2015) and to predict negatively biased learning about own abilities (Müller-Pinzler et al., 2022). Inversely, the experience of authentic pride, an emotion of accomplishment, is closely linked to the feeling of self-competence and self-worth (Stolz et al., 2020; Tracy et al., 2009), and predicts positively biased ability-belief formation (Müller-Pinzler et al., 2022).

From a clinical perspective, fostering self-compassion may be especially helpful for treating mental disorders that involve strong self-criticism, diminished self-efficacy, and low self-esteem (e.g., depression, anxiety). In line with this, low levels of self-compassion are seen as a risk factor for recurrent depressive episodes (Ehret et al., 2015) and correlate with depressive symptom burden (Krieger et al., 2013). Incorporating self-compassion practices into therapy may help individuals cope better with triggers such as negative feedback and the associated negative affective states, reduce rumination and self-criticism, and boost positive affect while maintaining a more stable self-esteem. Indeed, self-compassion interventions seem to offer effective treatment options for depression, anxiety and stress (Neff & Germer, 2013; Wilson et al., 2019). The present findings show that not self-compassion alone but also experienced affective states shape self-esteem. In major depression, it is assumed that negative self-images are maintained through heightened affective responsivity to negative feedback, devaluation of positive feedback and decreased associated positive affect (Czekalla et al., 2024). Similarly, other affective disorders such as social anxiety are characterized by generally lower positive and higher negative affect, which could entail inaccurate and overly pessimistic performance evaluation (Alden et al., 2008; Glazier & Alden, 2019). Thus, a better understanding of how affective experiences contribute to more global self-beliefs, possibly moderated by self-compassion, remains an important research topic.

## Conclusion

Our study demonstrates how self-compassion, positive affective experience, and subjective evaluations of performance feedback jointly contribute to changes in self-esteem. Self-compassion had a triple role: it predicted higher overall positive affect during the task, yielded higher self-esteem after the task, and protected the self-esteem of those individuals who showed low performance.

Positive affect, which was further driven by momentary expectations and experiences of success and failure during task execution, predicted self-esteem. Last, changes in self-esteem were predicted not by objective performance levels but rather by their subjective evaluation, possibly linked to implicit expectations of overall task performance and violations of these expectations. These findings are among the first to assess experimentally how self-compassion is tied to affective experience and self-evaluations. Further, they expand our understanding of the complex determinants of self-esteem, and offer therapeutically relevant perspectives on protecting and establishing self-esteem.

## Data and Code availability

The behavioral data and code used for analyses is available at https://osf.io/nd359/ (https://doi.org/10.17605/OSF.IO/ND359). The RMarkdown documents contain information about the structure and output of the code.

## Supporting information

Contains all supplementary materials

## Acknowledgments

We would like to thank the research assistants of our lab for their help in collecting the data for the present study. We also thank Nora Czekalla for sharing her expertise on self-compassion in psychotherapeutic practice. The research was supported by Else Kröner-Fresenius-Stiftung.

## Author Contributions

D.S.S., F.M.P. and S.K. conceived and designed the experiment. D.S. collected the data. J.F.W. analyzed the data. J.F.W., D.S.S. and F.M.P. discussed the data analysis and interpretation of results. The original draft of the manuscript was written by J.F.W. and D.S.S.. J.F.W., D.S.S., F.M.P., and S.K. reviewed and edited the paper.

