## Supplementary material for "Self-compassion is linked to positive affect and buffers declines in self-esteem in low performers": Contains all supplementary materials

### Supplementary figures

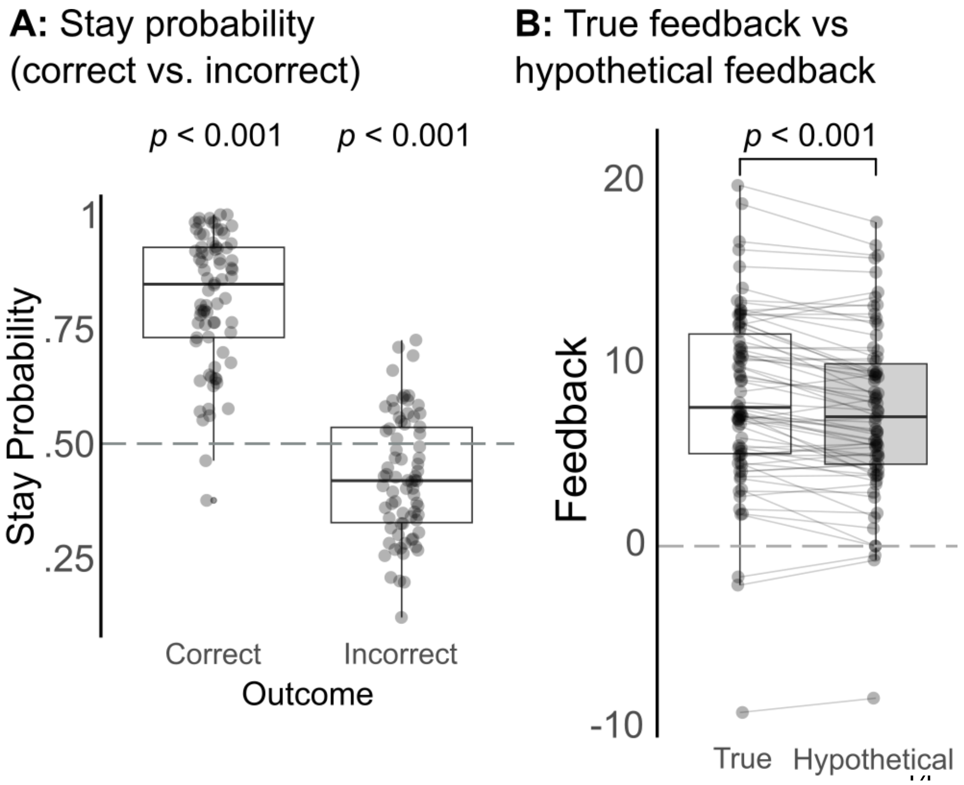


**Figure S1.** Participants adapt their behavior in response to task outcomes. **A)** Following correct choices, participants retained their choice significantly more often than chance (50%) (*median*=0.84, *IQR* 0.19) (*W*=2767, $p<0.001$), but significantly less often, following incorrect choices (*mean*=0.43, *sd*=0.13, $t(73)=-4.7$, $p<0.001$). **B)** Participants' feedback exceeded the hypothetical feedback they would have received if they had exerted the same amount of effort on each trial, irrespective of having chosen correctly or not ($t(73)=6.03, p<.001$).

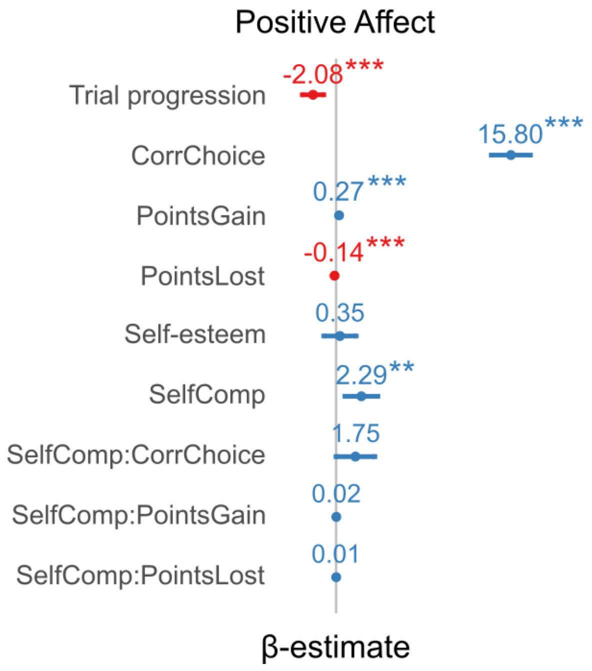


**Figure S2.** Task feedback is the main driver of positive affect. Running a linear mixed-model using task feedback variables (*conditional* $R^2$=0.72; *marginal* $R^2$=0.54) revealed that choosing the correct button ($\beta$=15.80, *s.e.*=0.89, $p$<0.001), gaining ($\beta$=0.27, *s.e.*=0.02, $p$<0.001) and losing points ($\beta$=-0.14, *s.e.*=0.02, $p$<0.001) significantly predicted positive affect during the task. Further, we found that self-compassion positively ($\beta$=2.29, *s.e.*=0.78, $p$=0.004) and task on time (trial progression, $\beta$=-2.08, *s.e.*=0.49, $p$<0.001) negatively predict positive affect. There was a trend towards an interaction between self-compassion and correct choice ($p$=0.055), whereas all other interactions remained non-significant (all *p*s>0.05). ** p < .01, *** p < .001

CorrChoice = correct choice; SelfComp = self-compassion

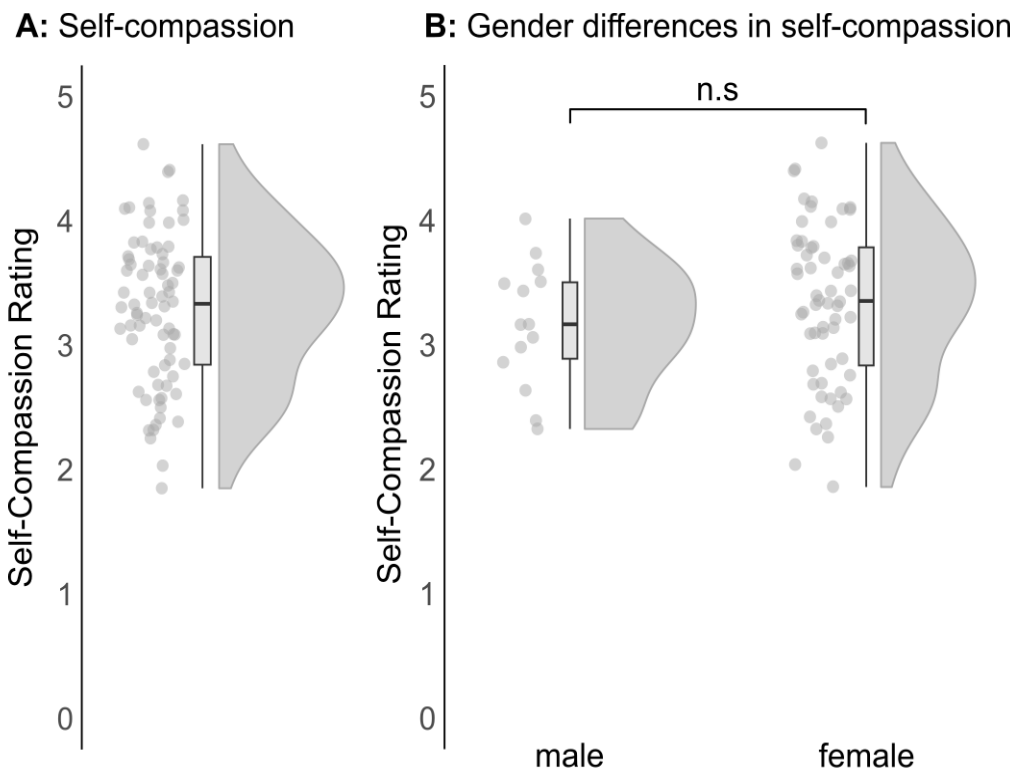


**Figure S3.** Self-compassion: descriptive statistics. **A)** Self-compassion was normally distributed (*mean*=3.29, *sd*=0.61, *W*=0.986, *p*=0.62). **B)** No gender differences were found (*t*(23)=1.03, *p*=0.32).

### Supplementary Tables

**Table S1.** Bivariate Pearson correlations and Cronbach's alpha. Strong positive correlation between positive affect, self-compassion and self-esteem. Correlation values are depicted in the center of each box, with 95% confidence intervals shown in brackets below. Internal consistency of items is plotted in the diagonal. SE = self-esteem; Pre = pre-task; Post = post-task

| Cronbach's α | SE (Pre) | SE (Post) | Self-Compassion | Positive Affect |
|---|---|---|---|---|
| SE (Pre) | **.85**<br>[.80, .90] | | | |
| SE (Post) | **.69**<br>[.55, .79] | **.87**<br>[.83, .91] | | |
| Self-Compassion | **.49**<br>[.29, .64] | **.52**<br>[.33, .67] | **.90**<br>[.87, .93] | |
| Positive Affect | **.24**<br>[.02, .45] | **.43**<br>[.22, .60] | **.36**<br>[.15, .55] | **.86**<br>[.81, .90] |

**Computational model**

*Hierarchical regression results for positive affect*

| Model | 0 | 1 | 2 | 3 | 4 | 5 | 6 | 7 | 8 | 9 | 10 | 11 | 12 | 13 |
|---|---|---|---|---|---|---|---|---|---|---|---|---|---|---|
| **Fixed effects** | | | | | | | | | | | | | | |
| Intercept | 48.93 | 53.33 | 49.20 | 51.72 | 52.02 | 52.00 | 51.98 | 52.17 | 52.17 | 52.17 | 52.16 | 52.16 | 52.16 | 52.16 |
| Trial progression | | -4.45 | 0.16 | -1.92 | -2.25 | -2.24 | -2.27 | -2.40 | -2.40 | -2.39 | -2.39 | -2.39 | -2.39 | -2.39 |
| RewExp | | | 12.24 | 8.90 | 7.98 | 8.01 | 7.92 | 7.74 | 7.77 | 7.77 | 7.78 | 7.78 | 7.79 | 7.79 |
| RewPE | | | | 11.03 | 11.04 | 11.04 | 10.98 | 11.08 | 11.09 | 11.09 | 11.09 | 11.09 | 11.09 | 11.09 |
| Ignored evidence | | | | | -1.36 | -1.34 | -1.38 | -1.53 | -1.51 | -1.52 | -1.51 | -1.51 | -1.51 | -1.51 |
| SE [Pre] | | | | | | | | | | 1.57 | 0.49 | 0.49 | 0.49 | 0.49 |
| SC | | | | | | | | | | | 2.25 | 2.25 | 2.29 | 2.35 |
| RewExp x SC | | | | | | | | | | | | 0.09 | 0.51 | 0.42 |
| RewPE x SC | | | | | | | | | | | | | 1.20 | 1.31 |
| Ign. evidence x SC | | | | | | | | | | | | | | -0.21 |
| **Random effects** | | | | | | | | | | | | | | |
| $\sigma_e^2$ | 492.3 | 485.5 | 342.7 | 222.4 | 221.3 | 217.8 | 191.3 | 153.5 | 151.2 | 151.2 | 151.2 | 151.2 | 151.2 | 151.2 |
| $\sigma_0^2$ | 32.7 | 32.8 | 34.9 | 36.2 | 36.2 | 33.6 | 29.5 | 32.4 | 33.7 | 30.2 | 25.7 | 25.7 | 25.6 | 25.6 |
| $\sigma_1^2$ | | | | | | 10.2 | 10.8 | 9.8 | 9.5 | 9.5 | 9.5 | 9.5 | 9.5 | 9.5 |
| $\sigma_2^2$ | | | | | | | 28.8 | 15.4 | 14.0 | 14.0 | 14.0 | 14.0 | 13.9 | 13.9 |
| $\sigma_3^2$ | | | | | | | | 38.6 | 38.8 | 38.8 | 38.8 | 38.8 | 37.1 | 37.1 |
| $\sigma_4^2$ | | | | | | | | | 4.0 | 4.0 | 4.0 | 4.0 | 4.0 | 4.0 |
| Deviance | 46939.4 | 46868.9 | 45090.2 | 42880.9 | 42856.2 | 42826.3 | 42335.5 | 41387.7 | 41356.0 | 41351.4 | 41342.6 | 41342.6 | 41339.8 | 41339.4 |

**Table S2.** Setup of computational hierarchical regression model. In Model 0-4, we iteratively added first-level regressors (e.g. performance information, trial progression) as fixed effects, in the order specified in the left-most column. Next, the respective random effects of these first-level regressors were added in Model 5-8. Last, we included the second-level predictors (e.g. self-compassion) and cross-level interactions between self-compassion and performance information in the order specified in the left-most column (Model 9-13).

**Behavioral model**

*Hierarchical regression results for positive affect*

| **Model** | 0 | 1 | 2 | 3 | 4 | 5 | 6 | 7 | 8 | 9 | 10 | 11 | 12 | 13 |
|---|---|---|---|---|---|---|---|---|---|---|---|---|---|---|
| **Fixed effects** | | | | | | | | | | | | | | |
| Intercept | 48.93 | 53.33 | 52.72 | 51.13 | 51.93 | 51.91 | 51.98 | 51.98 | 51.83 | 51.83 | 51.83 | 51.83 | 51.84 | 51.84 |
| Trial progression | | -4.45 | -3.04 | -1.41 | -2.17 | -2.14 | -2.21 | -2.22 | -2.08 | -2.08 | -2.08 | -2.08 | -2.08 | -2.08 |
| Correct Choice | | | 15.86 | 15.92 | 15.82 | 15.80 | 15.79 | 15.79 | 15.80 | 15.80 | 15.80 | 15.80 | 15.80 | 15.80 |
| Points gained | | | | 0.26 | 0.25 | 0.26 | 0.25 | 0.27 | 0.27 | 0.27 | 0.27 | 0.27 | 0.27 | 0.27 |
| Points lost | | | | | -0.16 | -0.16 | -0.14 | -0.13 | -0.14 | -0.14 | -0.14 | -0.14 | -0.14 | -0.14 |
| SE [Pre] | | | | | | | | | | 1.34 | 0.35 | 0.34 | 0.35 | 0.35 |
| SC | | | | | | | | | | | 2.17 | 2.30 | 2.24 | 2.29 |
| Choice x SC | | | | | | | | | | | | 1.59 | 1.81 | 1.75 |
| Gained x SC | | | | | | | | | | | | | 0.02 | 0.02 |
| Lost x SC | | | | | | | | | | | | | | 0.01 |
| **Random effects** | | | | | | | | | | | | | | |
| $\sigma_e^2$ | 492.3 | 485.5 | 229.6 | 218.3 | 214.8 | 211.5 | 149.7 | 146.9 | 142.8 | 142.8 | 142.8 | 142.8 | 142.8 | 142.8 |
| $\sigma_0^2$ | 32.7 | 32.8 | 36.5 | 36.8 | 36.7 | 37.1 | 38.4 | 39.2 | 37.1 | 35.2 | 30.1 | 30.1 | 30.1 | 30.1 |
| $\sigma_1^2$ | | | | | | 9.7 | 9.9 | 9.8 | 10.6 | 10.6 | 10.6 | 10.6 | 10.6 | 10.6 |
| $\sigma_2^2$ | | | | | | | 60.3 | 60.7 | 60.7 | 60.6 | 60.6 | 57.7 | 57.7 | 57.8 |
| $\sigma_3^2$ | | | | | | | | 0.02 | 0.02 | 0.02 | 0.02 | 0.02 | 0.02 | 0.02 |
| $\sigma_4^2$ | | | | | | | | | 0.03 | 0.03 | 0.03 | 0.03 | 0.03 | 0.03 |
| Deviance | 46939.4 | 46868.9 | 43044.5 | 42768.8 | 42704.4 | 42677.3 | 41175.9 | 41142.4 | 41058.0 | 41054.6 | 41046.9 | 41043.7 | 41042.0 | 41041.8 |

**Table S3.** Setup of behavioral hierarchical regression model. In Model 0-4, we iteratively added first-level regressors (e.g. task feedback, trial progression) as fixed effects, in the order specified in the left-most column. Next, the respective random effects of these first-level regressors were added in Model 5-8. Last, we added the second-level predictors (e.g. self-compassion) and cross-level interactions between self-compassion and task feedback in the order specified in the left-most column (Model 9-13).
